# A murine model to study chronic airway fungal colonisation that recapitulates human disease

**DOI:** 10.64898/2026.05.20.726561

**Authors:** Emily A. Sey, Hilaire Irere, Adilia Warris, Fabián Salazar

**Affiliations:** MRC Centre for Medical Mycology at the University of Exeter, Department of Biosciences, Exeter, United Kingdom

## Abstract

*Aspergillus fumigatus* is a ubiquitous environmental mould and a leading cause of chronic fungal-associated respiratory disease, yet the mechanisms by which persistent airway colonisation drives immune adaptation and lung pathology remain poorly understood. Progress in this area has been limited by the lack of *in vivo* models that recapitulate stable, non-invasive fungal persistence without immunosuppression. Here, we developed and optimised a murine model of chronic airway colonisation using agar bead–embedded *A. fumigatus* conidia delivered intratracheally. Embedding did not impair fungal germination or hyphal growth, and the agar matrix was immunologically inert, supporting its use as a neutral scaffold. This approach established stable fungal persistence in the airways for at least three weeks in immunocompetent mice without inducing invasive disease or systemic morbidity. Colonisation elicited a transient, airway-restricted innate immune response characterised by early neutrophil and monocyte recruitment and increased CXCL1, MIP-1α, MIP-1β, and TNF production, which resolved over time. Histopathological analysis revealed a progressive sequence of disease-relevant features, including initial immune containment, followed by mucus hypersecretion, and airway remodelling. At the adaptive level, persistent colonisation induced a dynamic T cell response that transitioned from an early polyfunctional profile to a sustained Th17-dominant phenotype. Importantly, application of this model in CFTR-deficient mice uncovered enhanced collagen deposition and fibrotic remodelling without altered fungal burden, demonstrating its utility in modelling disease-relevant outcomes in susceptible hosts. Together, this study establishes a robust and physiologically relevant platform for investigating host–fungal interactions during chronic airway colonisation. This model provides new opportunities to dissect mechanisms of immune adaptation, fungal persistence, and tissue remodelling, and to identify therapeutic strategies targeting chronic *Aspergillus*-associated lung disease.

## Introduction

*Aspergillus fumigatus* is a ubiquitous environmental mould and a leading cause of fungal-associated respiratory disease. Recently designated a WHO “critical priority pathogen,” *A. fumigatus* poses a substantial threat to global health^1^. Humans are continuously exposed to airborne conidia, and although immunocompetent individuals typically clear these efficiently, people with impaired immunity or structural lung abnormalities are highly vulnerable to persistent colonisation^2^. Indeed, up to 60% of individuals with cystic fibrosis, 30% with COPD, and 25% with bronchiectasis exhibit airway colonisation by *Aspergillus*^3-8^. Fungal sensitisation is common in asthma and contributes significantly to morbidity and mortality^9-12^. Despite this, the biological mechanisms through which *A. fumigatus* shapes chronic lung inflammation and disease progression remain poorly defined.

*A. fumigatus* causes a spectrum of clinical syndromes ranging from allergic airway disease to chronic pulmonary aspergillosis (CPA) and invasive infection^13^. While considerable progress has been made in defining the immune mechanisms that prevent invasive aspergillosis, far less is known about host responses to persistent fungal exposure or non-invasive colonisation^14^. A major limitation has been the lack of tractable *in vivo* models that reproduce stable fungal persistence without relying on immunosuppression. Current models typically induce invasive infection or fail to capture the airway-limited, neutrophil-driven inflammation characteristic of CPA and *Aspergillus* bronchitis. Moreover, defining and studying colonisation in patients is challenging due to its dynamic and often transient nature^15,16^.

To address these gaps, we developed and optimised a murine model in which *A. fumigatus* conidia are embedded in bio-inert agar beads and delivered intratracheally. Agar-bead microenvironments mimic aspects of the airway matrix, enabling stable fungal persistence and sustained host–pathogen interactions without causing invasive disease^17-19^. This methodology, commonly used in chronic bacterial infection models^20^, has not been widely applied in fungal research.

Here, we show that this approach establishes reliable, non-invasive colonisation for at least three weeks, accompanied by airway-restricted inflammation that recapitulates key features of human *Aspergillus*-associated disease. By day 7, colonised mice exhibit marked neutrophil and monocyte recruitment, along with increased production of CXCL1 and MIP-1α. By day 14, this progresses to focal mucus accumulation and airway remodelling, consistent with features of CPA and *Aspergillus* bronchitis. At the adaptive level, persistent colonisation induces a dynamic T cell response that transitions from an early polyfunctional profile to a sustained Th17-dominant programme, suggesting a key role for Th17 immunity in controlling fungal persistence while limiting excessive inflammation. In a cystic fibrosis model, colonisation further exacerbates lung pathology, promoting increased fibrosis without affecting fungal burden, and highlighting its contribution to disease progression in susceptible hosts.

Together, this work establishes a robust and physiologically relevant platform for dissecting the immunological, epithelial, and structural drivers of chronic *Aspergillus* colonisation. This model provides an essential tool for advancing mechanistic insights into fungal persistence, immune dysregulation, and disease progression in chronic respiratory disorders.

## Results

### Embedding in agar beads does not alter *Aspergillus fumigatus* growth kinetics

To establish a chronic lung colonisation model using agar bead–embedded *Aspergillus* conidia, we first assessed whether embedding affected *A. fumigatus* germination and hyphal growth compared with free conidia. The germination rate at 10 hours did not differ significantly between free conidia and conidia embedded in agar beads (Fig. 1A-B). We next evaluated hyphal outgrowth and found that hyphal length was comparable between conditions up to 8 hours. At this time point, conidia embedded in agar beads exhibited a modest but significant increase in hyphal length compared with free conidia (mean length 35.30 µm vs 23.60 µm, respectively; Fig. 1C). By 10 hours, however, hyphal length was no longer significantly different between free and embedded conidia (mean length 43.90 µm vs 54.19 µm, respectively; Fig. 1C). Together, these data demonstrate that embedding *A. fumigatus* conidia in agar beads does not impair fungal germination or hyphal growth, supporting the use of this approach to model chronic airway colonisation.

**Fig. 1.**
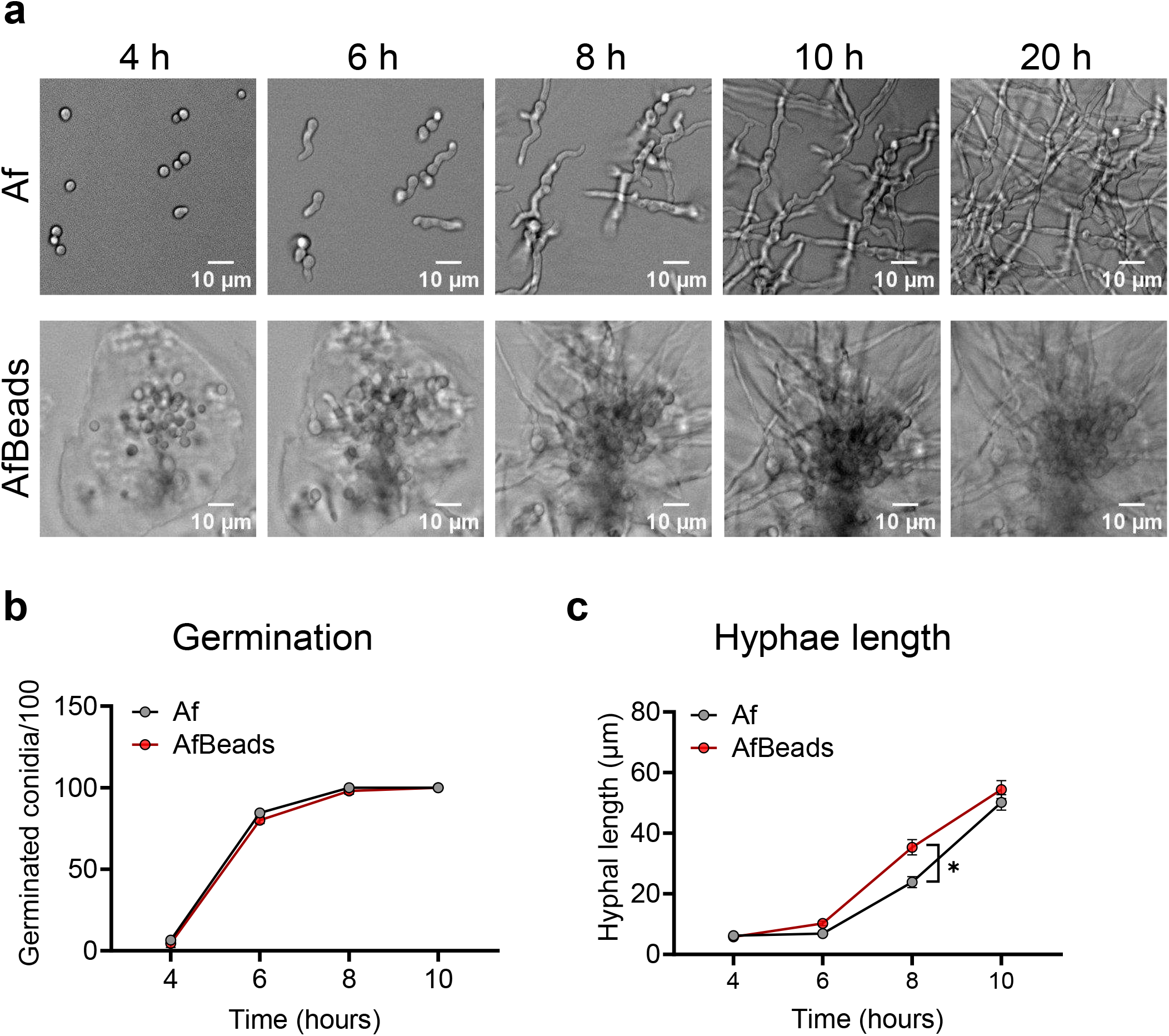
Embedding in agar beads does not alter Aspergillus fumigatus growth kinetics. **(a)** Free A. fumigatus conidia (top) or conidia embedded in agar beads (bottom) were incubated in medium, and representative digital images (20× magnification) were acquired at 4, 6, 8, and 10 hours to assess fungal growth and morphogenesis. **(b–c)** Free conidia (black line) or agar bead–embedded conidia (red line) were cultured in medium on µ-Slide 8-well chambered coverslips. Time-lapse images (1024 × 1024 pixels, CZI format) were acquired, and 100 individual conidia per condition were tracked. Germination rates **(b)** and hyphal length **(c)** were quantified at 4, 6, 8, and 10 hours using ZEISS ZEN 3.8 software. Each point represent mean±SEM of pooled data from two independent experiments. Statistical analysis was performed using Two-way ANOVA; *p < 0.05.

### Empty agar beads do not induce epithelial cell activation

We next determined whether agar beads themselves could induce inflammatory responses that might confound interpretation of host responses to bead-embedded *Aspergillus*. To test this, we examined the effect of sterile, empty agar beads on airway epithelial cells in vitro. Human immortalised bronchial epithelial cells (HBEC) were exposed to empty agar beads for 24 hours, and secretion of the pro-inflammatory cytokines IL-8 and IL-1β was measured. Empty agar beads did not induce significant cytokine production compared with unstimulated controls, indicating that the beads alone do not trigger epithelial inflammatory responses (Fig. 2A). Similar results were obtained using human immortalised cystic fibrosis bronchial epithelial cells (CFBE41o–) (Fig. 2B). Together, these findings demonstrate that the agar bead matrix is immunologically inert in this epithelial system and therefore represents a suitable neutral scaffold for modelling persistent fungal colonisation

**Fig. 2.**
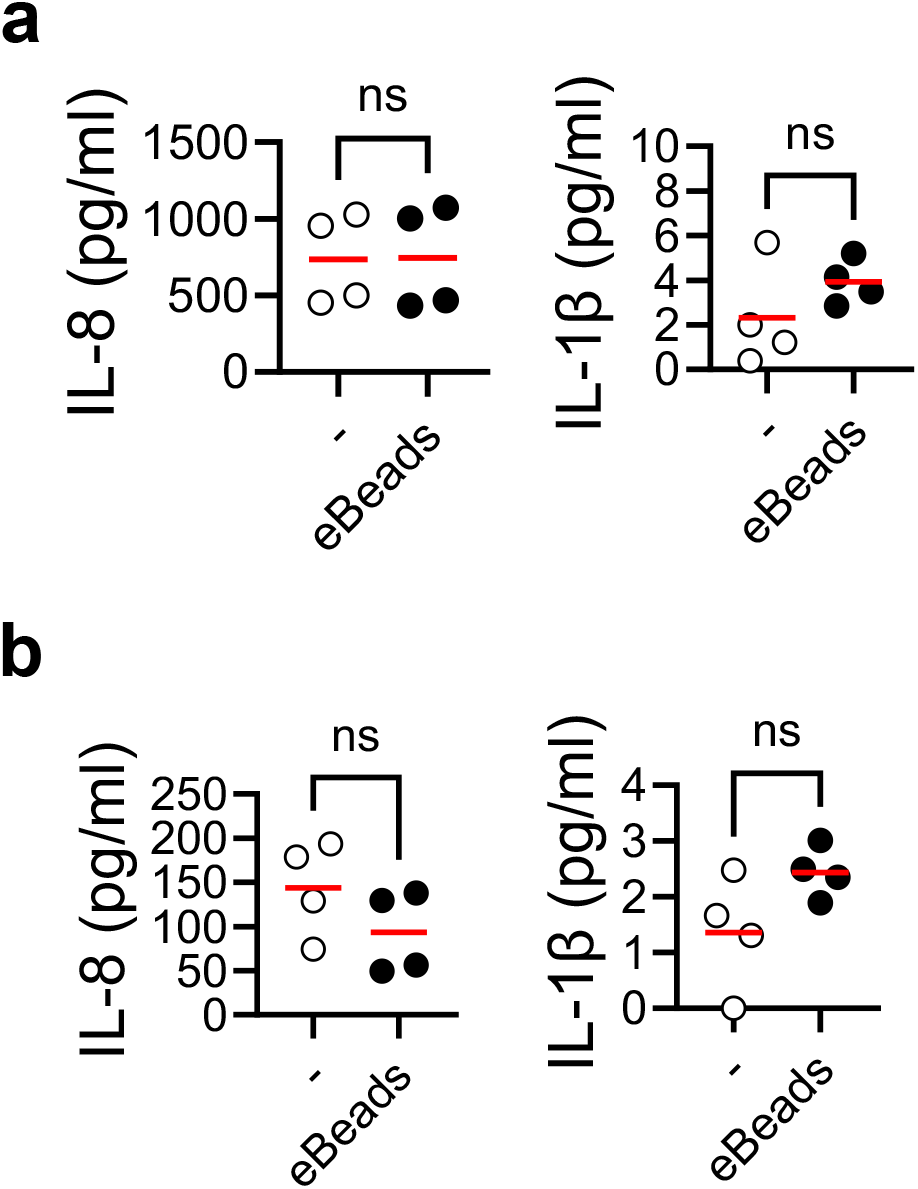
Agar beads do not induce epithelial cell activation. Human immortalised bronchial epithelial cells (HBEC) (a) and human immortalised CF bronchial epithelial cells (CFBE41o–) (b) were stimulated with sterile, empty agar beads or left unstimulated as a control. After 24 hours, cell culture supernatants were collected, and IL-8, and IL-1β levels measured by ELISA. Red line represent mean of pooled data from two independent experiments. Statistical analysis was performed using Student’s t-test; *p < 0.05.

### Agar bead–embedded *Aspergillus* establishes stable pulmonary colonisation

To determine whether agar bead-embedded *Aspergillus* conidia can establish sustained airway colonisation *in vivo, A. fumigatus* spores embedded in agar beads were delivered intratracheally into mice and fungal persistence was monitored for up to three weeks (Fig. 3A). Throughout the observation period, mice exhibited no overt clinical signs of disease. A transient and modest reduction in body weight was observed at day 2 post-challenge, which fully resolved by day 5 (Fig. 3B), indicating minimal systemic morbidity associated with fungal colonisation. Assessment of lung fungal burden revealed that colony-forming units (CFU) recovered from lung tissue remained relatively stable at days 7, 14, and 21 post-challenge (Fig. 3C). The absence of progressive fungal clearance or uncontrolled fungal expansion suggests that agar bead embedding enables *A. fumigatus* to persist within the lung in a controlled, non-invasive manner. Together, these data demonstrate that agar bead–embedded *Aspergillus* establishes a stable state of pulmonary colonisation that is maintained over several weeks without inducing acute disease.

**Fig. 3.**
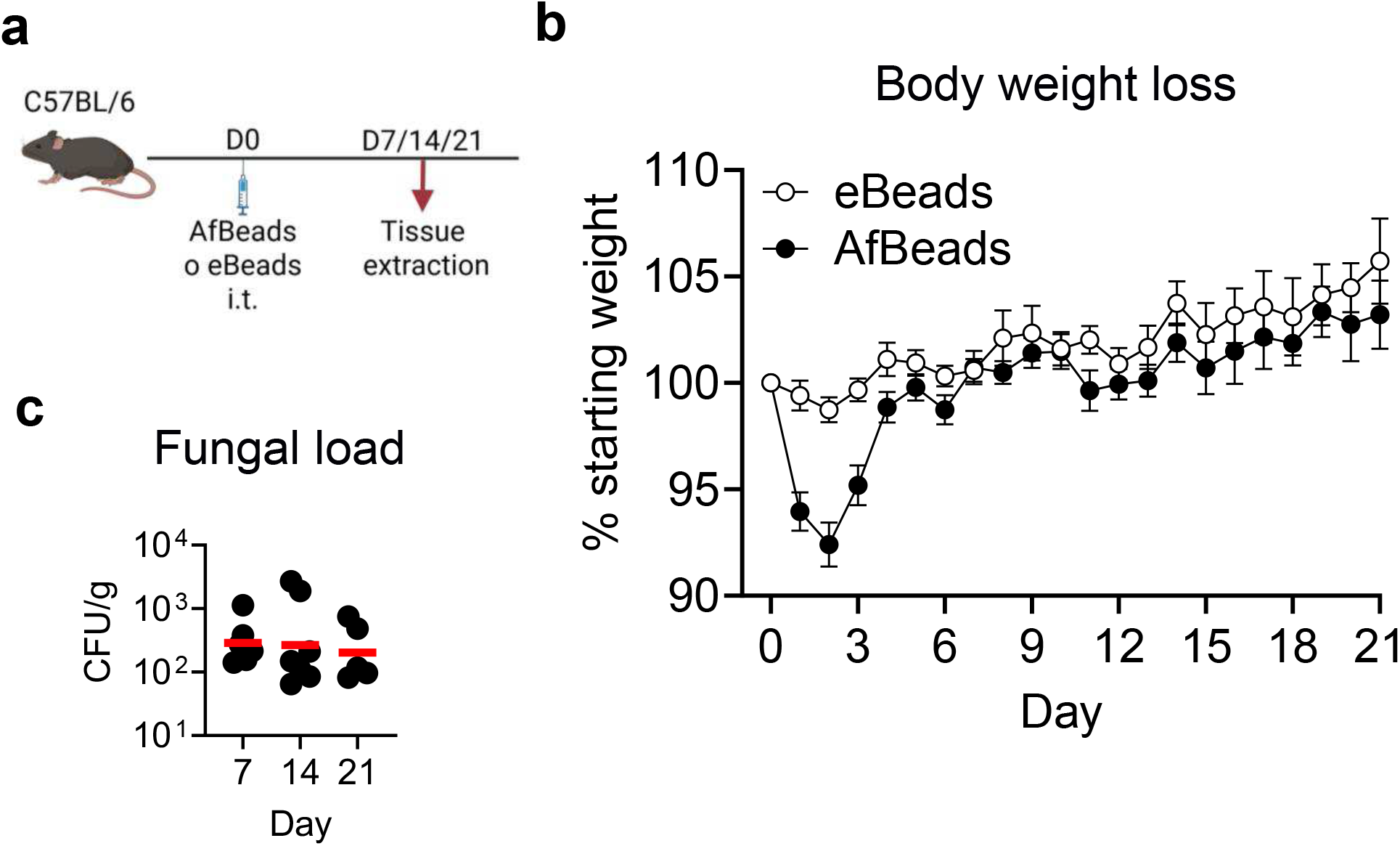
Agar bead–embedded Aspergillus establishes stable airway colonisation. A. fumigatus conidia were embedded in agarose beads and delivered intratracheally into C57BL/C mice. (a) Schematic overview of the agar bead colonisation model. (b) Changes in body weight monitored over a 3-week period following challenge. Each point represent mean±SEM of pooled data from two independent experiments. (c) Lung fungal burden quantified at days 7, 14, and 21 post-challenge and expressed as colony forming units (CFUs) per gram of lung tissue. Each point represents an individual mouse. Red line represent mean of pooled data from two independent experiments.

### Agar bead–embedded *Aspergillus* induces histopathological features resembling human disease

We next assessed whether agar bead–embedded *Aspergillus* colonisation recapitulates key histopathological features observed in human airway disease. Lung tissue was analysed by periodic acid–Schiff (PAS) staining at days 7, 14, and 21 post-challenge. At day 7, pronounced inflammatory cell infiltration was observed surrounding agar beads within the airways, with fungal hyphae clearly visible but largely confined within the bead matrix (Fig. 4A). This early response suggests effective immune containment of fungal growth without evidence of tissue invasion. By day 14 post-challenge, inflammatory infiltrates were markedly reduced, coinciding with the emergence of fungal hyphae extending beyond the beads and establishing direct interactions with the airway epithelium. This stage was associated with a noticeable increase in mucus deposition within the airways, consistent with epithelial activation and goblet cell responses. By day 21, histological analysis revealed features of airway remodelling, including altered airway architecture, indicative of sustained tissue adaptation to chronic fungal presence (Fig. 4A). Importantly, mice challenged with sterile, empty agar beads did not exhibit detectable airway inflammation, mucus accumulation, or structural remodelling, confirming that these pathological changes are driven by fungal colonisation rather than the delivery matrix itself. Quantitative analysis of mucus deposition at day 14 demonstrated a significant increase in airway mucus area in *Aspergillus*-challenged mice compared with empty bead controls (Fig. 4B). Collectively, these findings demonstrate that agar bead–embedded *Aspergillus* induces a progressive sequence of histopathological changes, spanning early immune containment, epithelial engagement, mucus hypersecretion, and airway remodelling, that closely resemble features of chronic human *Aspergillus*-associated airway disease.

**Fig. 4.**
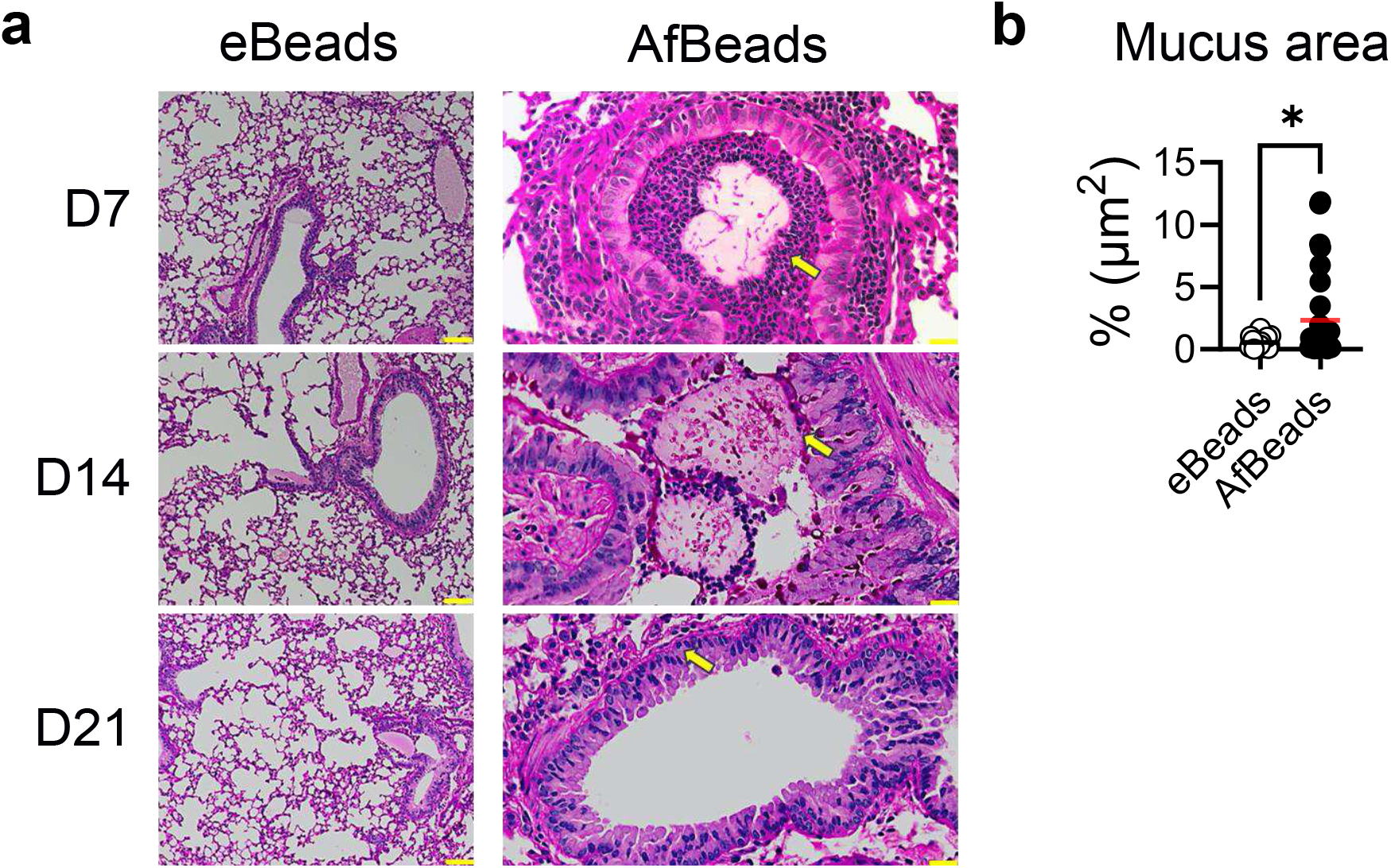
Agar bead-embedded Aspergillus induces histopathological features resembling human disease. A. fumigatus conidia embedded in agarose beads were delivered intratracheally into C57BL/C mice. (a) Representative lung sections stained with periodic acid–Schiff (PAS) at days 7, 14, and 21 post-challenge, highlighting cellular infiltration, fungal material and associated mucus production (yellow arrows). (b) Quantification of mucus plug area in lung sections at day 14 post-challenge, measured using QuPath software. Red line represent mean of pooled data from two independent experiments. Statistical analysis was performed using Student’s t-test; *p < 0.05.

### Agar bead–embedded *Aspergillus* induces a transient airway inflammatory response

To characterise the cellular inflammatory response elicited by agar bead–embedded *Aspergillus* colonisation, we quantified immune cell infiltration in the bronchoalveolar lavage fluid (BALF) and lung tissue at defined time points following intratracheal challenge. At day 7 post-challenge, we observed a significant increase in neutrophils and a trend towards increased inflammatory monocytes in the BALF compared to empty bead controls (Fig. 5A-B), indicative of an acute innate inflammatory response to fungal presence. Notably, this increase was transient, with neutrophil and monocyte numbers returning to baseline levels by day 14. In contrast, no significant changes were detected in eosinophils, dendritic cells, or alveolar macrophages in the BALF across the time points examined, suggesting the absence of a sustained type 2 response or broad disruption of resident myeloid populations (Fig. 5A-B). Consistent with these findings, analysis of immune cell populations within lung tissue revealed no major differences between *Aspergillus*-challenged mice and empty bead controls at any time point (Fig. 5C-D), indicating that inflammation was largely confined to the airway lumen rather than the lung parenchyma. Together, these data demonstrate that agar bead–embedded *Aspergillus* induces a localised and self-resolving airway inflammatory response, characterised by transient recruitment of innate immune cells without progression to chronic or invasive lung inflammation. This pattern closely mirrors key features of controlled fungal colonisation in the airways and further supports the suitability of this model for studying host responses to persistent, non-invasive *Aspergillus* exposure.

**Fig. 5.**
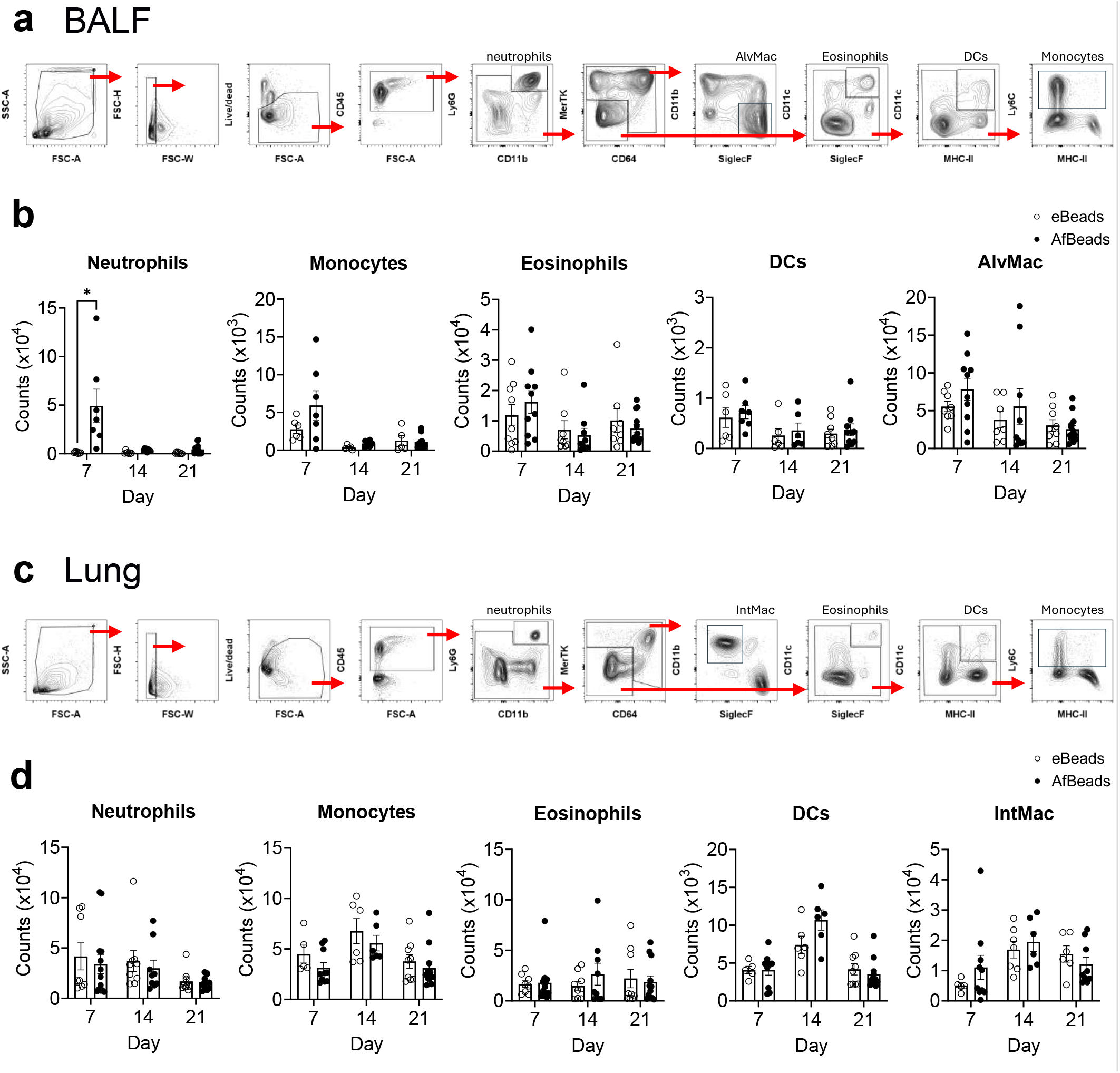
Agar bead–embedded Aspergillus induces a transient airway inflammatory response. A. fumigatus conidia embedded in agarose beads were delivered intratracheally into C57BL/6 mice. (a) Immune cell infiltration in bronchoalveolar lavage fluid (BALF) at days 7, 14, and 21 post-challenge, quantified by flow cytometry. (b) Immune cell infiltration in lung tissue at days 7, 14, and 21 post-challenge, quantified by flow cytometry. Each point represents an individual mouse. Bars represent mean±SEM of pooled data from two independent experiments. Statistical analysis was performed using Two-way ANOVA; *p < 0.05.

### Agar bead–embedded *Aspergillus* induces a transient airway cytokine response

We next characterised the airway cytokine milieu following challenge with agar bead–embedded *Aspergillus*. Analysis of BALF revealed a transient increase in pro-inflammatory cytokines and chemokines at day 7 post-challenge, including CXCL1, MIP-1α, MIP-1β, and TNF (Fig. 6). These mediators are associated with neutrophil recruitment and early innate immune activation, consistent with the transient cellular influx observed in the airways at this time point. By contrast, no significant changes were detected in a broad panel of additional cytokines and chemokines at any time point examined, including MCP-1, RANTES, eotaxin, IL-12p40, IL-12p70, IL-6, IL-10, G-CSF, IFN-γ, IL-2, IL-4, IL-5, or IL-17A. The absence of sustained pro-inflammatory, type 1, type 2, or type 17 cytokine signatures indicates that the immune response does not progress to chronic inflammation or overt immunopathology. Together, these data demonstrate that agar bead–embedded *Aspergillus* elicits a restricted and self-limiting airway cytokine response, characterised by early innate immune activation that rapidly resolves. This cytokine profile is consistent with controlled fungal colonisation not progressing to invasive infection and supports the use of this model to study immune adaptation and tolerance to persistent *Aspergillus* exposure in the airway.

**Fig. 6.**
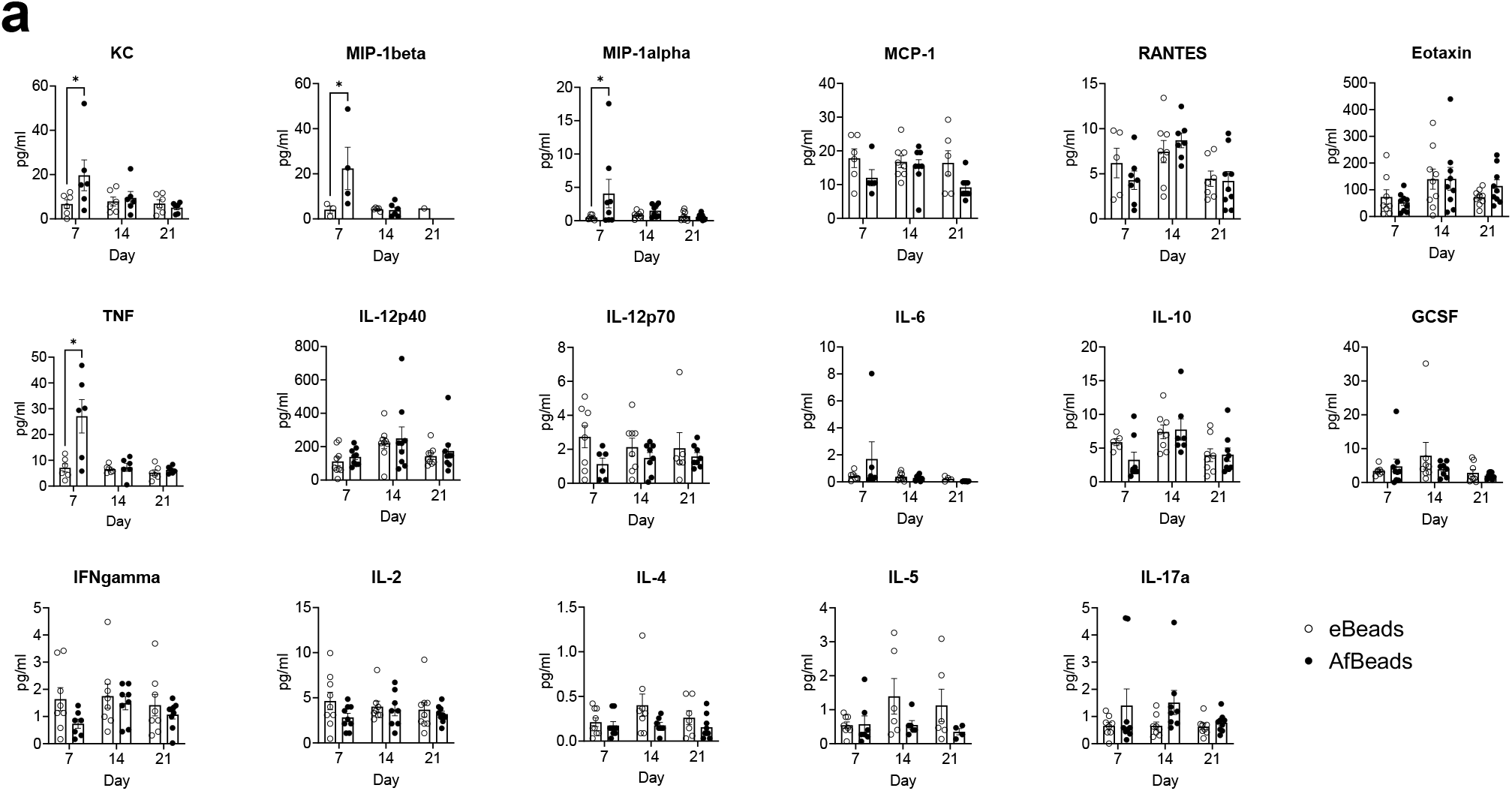
Agar bead–embedded Aspergillus induces a transient airway cytokine response. A. fumigatus conidia embedded in agarose beads were delivered intratracheally into C57BL/C mice. (a) Cytokine levels in bronchoalveolar lavage fluid (BALF) were quantified by Luminex analysis at days 7, 14, and 21 post-challenge. Each point represents an individual mouse. Bars represent mean±SEM of pooled data from two independent experiments. Statistical analysis was performed using Two-way ANOVA; *p < 0.05.

### Agar bead–embedded *Aspergillus* elicits a Th17-dominant pulmonary T cell response

To define the nature of the adaptive T cell response during chronic *Aspergillus* colonisation, we assessed pulmonary T cell phenotypes using an *in vitro* recall assay at defined time points following intratracheal challenge with agar bead–embedded *A. fumigatus* (Fig. 7A). At day 7 post-challenge, we observed a significant increase in cytokine-producing T cells compared with empty bead controls. This early response was characterised by a polyfunctional T cell profile, with increased production of IFN-γ, IL-10, IL-17A, and IL-4, indicating the simultaneous engagement of Th1-, Th2-, Th17-, and regulatory-associated pathways during the initial phase of fungal colonisation (Fig. 7B). Strikingly, by day 21 post-challenge, the T cell response had evolved to a more restricted phenotype, with IL-17A emerging as the dominant cytokine. At this later time point, no significant increases in IFN-γ, IL-4, or IL-10 production were detected relative to controls, indicating a selective maintenance of Th17 responses during fungal persistence (Fig. 7C). Together, these data demonstrate that agar bead–embedded *Aspergillus* drives a dynamic and temporally regulated adaptive immune response, transitioning from an early, broadly polyfunctional T cell activation to a Th17-dominant pulmonary T cell phenotype during chronic colonisation. This pattern is consistent with a role for Th17 immunity in maintaining fungal containment at the airway surface while avoiding excessive inflammation, and closely mirrors adaptive immune signatures observed in chronic *Aspergillus*-associated airway disease.

**Fig. 7.**
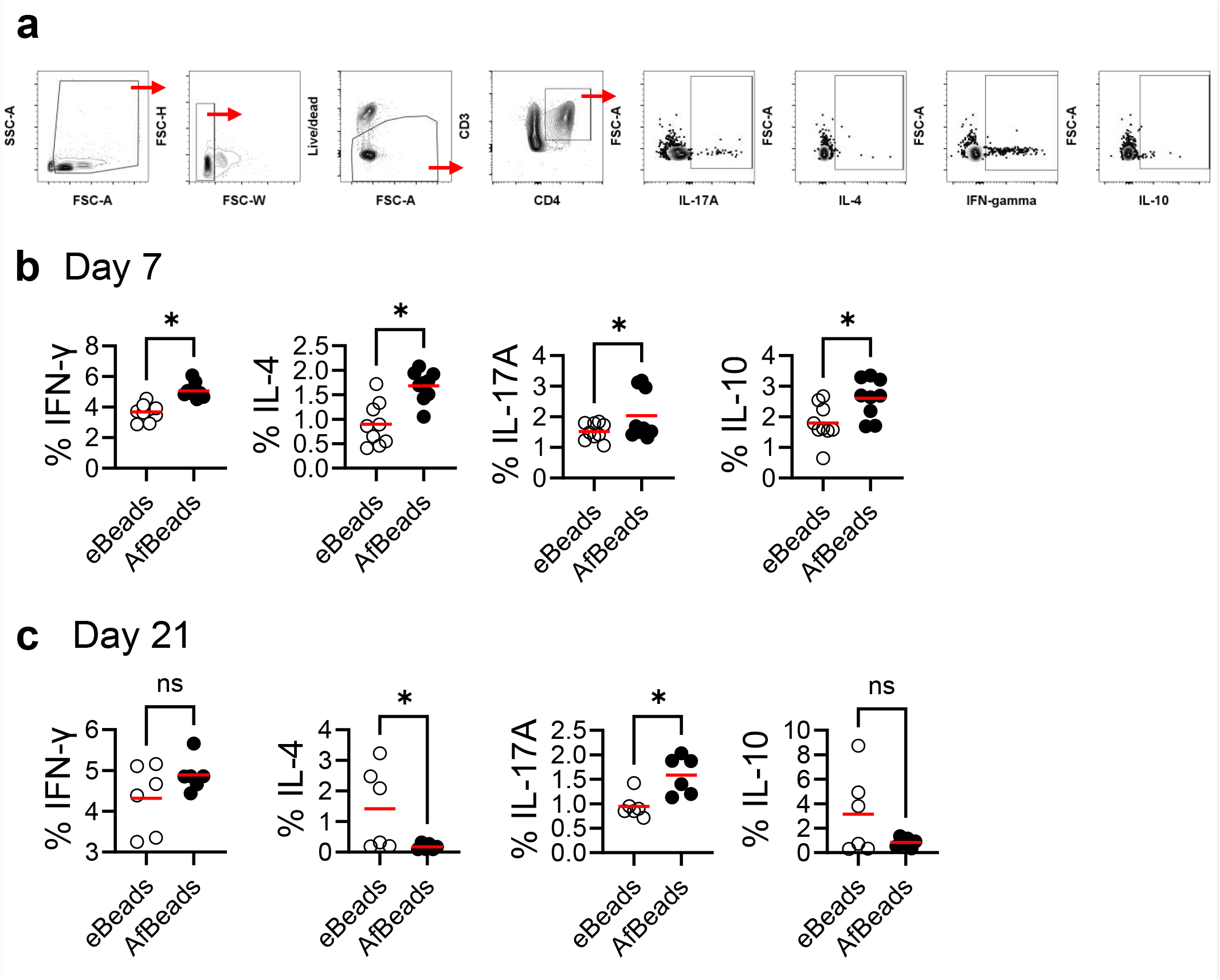
Agar bead–embedded Aspergillus elicits a Th17-dominant pulmonary T cell response. A. fumigatus conidia embedded in agarose beads were delivered intratracheally into C57BL/6 mice. (a) Intracellular cytokine staining of lung T cells at day 7 post-challenge following an in vitro whole-lung recall assay. (b) Intracellular cytokine staining of lung T cells at day 21 post-challenge following an in vitro whole-lung recall assay. Each point represents an individual mouse. Red line represent mean of pooled data from two independent experiments. Statistical analysis was performed using Student’s t-test; *p < 0.05.

### Colonisation of CFTR-deficient mice with agar bead–embedded *Aspergillus* exacerbates lung fibrosis

Finally, to assess the relevance of this colonisation model in the context of chronic lung disease, we applied the agar bead–embedded *Aspergillus* challenge to cystic fibrosis transmembrane conductance regulator–deficient (CFTR KO) mice (Fig. 8A). CFTR KO mice represent a well-established experimental model of cystic fibrosis-like lung disease, recapitulating the absence of functional CFTR in the airway epithelium and associated defects in mucociliary clearance and tissue homeostasis^21^. At day 14 post-challenge, no differences were observed in lung fungal burden (Fig. 8B), while histological analysis revealed a significant increase in collagen deposition in the lungs of CFTR KO mice compared with CFTR heterozygous control mice, as quantified by Sirius Red staining (Fig. 8C-D). This enhanced collagen accumulation indicates increased fibrotic remodelling in response to persistent *Aspergillus* colonisation in the absence of functional CFTR. These findings demonstrate that agar bead–embedded *Aspergillus* colonisation unmasks disease-relevant pathological outcomes in a genetically susceptible host, driving exaggerated fibrotic responses in CFTR-deficient mice. Collectively, this supports the translational utility of this model for investigating how chronic fungal colonisation contributes to airway remodelling and fibrosis in cystic fibrosis and other chronic lung diseases.

**Fig. 8.**
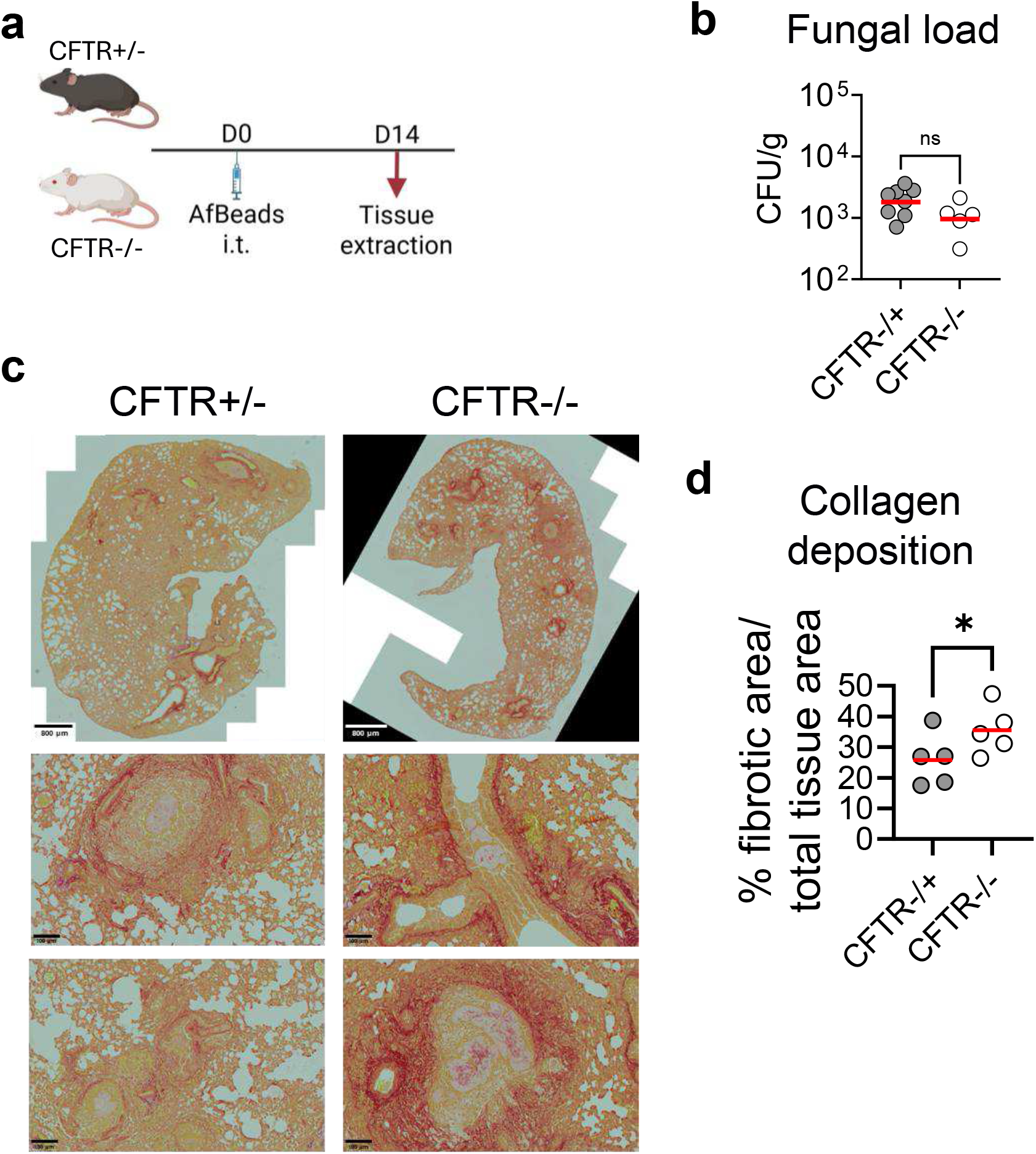
Colonisation of CFTR-deficient mice with agar bead–embedded Aspergillus exacerbates lung fibrosis. A. fumigatus conidia embedded in agarose beads were delivered intratracheally into CFTR+/- and CFTR-/-mice. (a) Schematic overview of the agar bead colonisation model. (b) Lung fungal burden quantified at day 14, post-challenge and expressed as colony forming units (CFUs) per gram of lung tissue. (c) Sirius red stain of lung histological sections (scale bar: 800 µm top, 100 µm bottom). (d) Quantification of fibrotic area in lung sections at day 14 post-challenge, measured using QuPath software. Each point represents an individual mouse. Red line represent mean of pooled data from one experiment. Statistical analysis was performed using Student’s t-test; *p < 0.05.

## Discussion

In this study, we describe and validate a murine model of chronic pulmonary *Aspergillus fumigatus* colonisation based on intratracheal delivery of agar bead–embedded conidia. This approach enables stable fungal persistence in the airway over several weeks without progression to invasive disease, while eliciting immune, epithelial, and tissue responses that closely resemble features of chronic *Aspergillus*-associated airway pathology in humans^22-24^. Our findings establish this model as a tractable experimental platform to interrogate host–fungal interactions during sustained airway colonisation rather than acute infection.

A central feature of this system is the use of agarose beads as a bio-inert scaffold that supports fungal persistence without directly perturbing host responses. We demonstrate that embedding *A. fumigatus* conidia in agar beads does not impair germination or hyphal growth kinetics in vitro, indicating that the matrix does not impose growth-restrictive or stress-inducing effects on the fungus. Importantly, agar beads themselves were immunologically inert in airway epithelial cultures, failing to induce pro-inflammatory cytokine production. Together, these observations confirm that host responses observed in vivo are driven by fungal persistence.

Beyond their inertness, agarose beads likely contribute biologically relevant physical cues. At the concentrations used, agarose exhibits stiffness and mechanical properties comparable to elements of the airway extracellular matrix (ECM), enabling host–pathogen interactions to occur within a confined yet permissive microenvironment^17-19^. Such physical constraint may facilitate prolonged fungal residence while limiting tissue invasion, mimicking conditions encountered in chronically diseased airways characterised by altered mucus composition, epithelial damage, and ECM remodelling. Future studies could build on this platform by exploring alternative matrices, including hydrogel-based or ECM-mimetic systems, which would allow finer control over mechanical stiffness, degradability, and biochemical signalling. This may provide further insight into how biophysical properties of the airway niche influence fungal growth, immune sensing, and tissue remodelling during chronic disease.

Following intratracheal delivery, agar bead–embedded *Aspergillus* established a stable pulmonary burden that was maintained for at least three weeks in immunocompetent mice, without weight loss, overt clinical disease, or uncontrolled fungal expansion. This distinguishes the model from conventional inhalation or intranasal challenge systems, which typically result in rapid fungal clearance in immunocompetent hosts or, alternatively, invasive disease when combined with immunosuppression. Instead, the stable CFU levels observed here more closely reflect the clinical scenario of airway colonisation seen in conditions such as cystic fibrosis (CF), chronic obstructive pulmonary disease, and severe asthma, where *Aspergillus* persists at the mucosal surface in the absence of angioinvasion^3-7^.

Histopathological analysis revealed a progressive sequence of tissue responses that mirror key features of human chronic airway disease. Early immune cell accumulation surrounding bead-localised fungi at day 7 suggested effective containment of fungal growth. Over time, this response transitioned towards epithelial engagement, mucus hypersecretion, and airway remodelling, with hyphae extending beyond the bead matrix by day 14 and structural changes evident by day 21. The absence of pathology in mice receiving empty beads confirms that these changes are fungus-driven. Notably, the temporal dissociation between early inflammation and later tissue remodelling mirrors human disease, where chronic structural changes often persist despite limited overt inflammation. Consistent with this, we observed a transient, airway-restricted innate immune response characterised by early recruitment of neutrophils and monocytes and a short-lived increase in pro-inflammatory mediators. This response resolves by day 14 without sustained cytokine production or changes in lung tissue immune populations, indicating controlled, non-invasive colonisation confined to the airway.

A particularly striking finding was the dynamic nature of the adaptive immune response. Early during colonisation, pulmonary T cells displayed a broad, polyfunctional cytokine profile encompassing Th1-, Th2-, Th17-, and regulatory-associated cytokines. Over time, this response consolidated into a Th17-dominant phenotype, with IL-17A emerging as the sole persistently elevated cytokine by day 21. Th17 immunity has been implicated in mucosal antifungal defence and fungal containment, but also in chronic airway pathology when dysregulated^25,26^. The emergence of a focused Th17 response in this model suggests a role for IL-17A in maintaining long-term fungal containment at the airway surface while avoiding excessive inflammation, a balance that is likely critical in chronic colonisation states.

Importantly, the absence of detectable IL-17A in bronchoalveolar lavage fluid despite the presence of Th17 cells in lung tissue highlights a key spatial and functional distinction. IL-17A production may be highly localised within the lung parenchyma or airway wall, acting in a cell-restricted manner that is not readily captured in lavage fluid. Alternatively, Th17 cells may be poised for rapid cytokine production upon antigen re-encounter rather than constitutively secreting IL-17A in vivo, as revealed by the recall assay. This dissociation between cellular potential and soluble cytokine detection underscores the importance of assessing tissue-resident immune responses rather than relying solely on airway fluid readouts when studying chronic colonisation.

Importantly, the translational relevance of this model was underscored by its application to CFTR-deficient mice. In this genetically susceptible host, agar bead–embedded *Aspergillus* colonisation resulted in significantly increased collagen deposition and fibrotic remodelling compared with heterozygous controls. These findings align with clinical observations linking *Aspergillus* colonisation to worse structural lung disease in CF^3,8^ and demonstrate that this model can unmask disease-relevant pathological consequences that are not apparent in immunocompetent hosts. The convergence of Th17 responses, and fibrosis in CFTR-deficient lungs highlights mechanistic pathways that warrant further investigation.

The agar bead model is well established for studying chronic bacterial lung infections^20^ and has been adapted for *Aspergillus* colonisation^27-30^. Compared to previous studies showing sustained neutrophilic inflammation^27^, our model provides a more comprehensive and longitudinal view, revealing a transient innate response followed by mucus accumulation and airway remodelling, better reflecting chronic human disease. These differences likely stem from the use of smaller beads and lower fungal doses, which may promote containment rather than prolonged inflammation. Additionally, differences in microbiome composition between animal facilities are known to influence experimental outcomes and contribute to variability between studies^31^, highlighting the need to further investigate how local microbial communities shape fungal colonisation and disease progression. While this study establishes a robust platform for modelling chronic *Aspergillus* colonisation, limitations remain. Agar bead embedding might represent an artificial constraint on fungal localisation, and future work will be required to determine how closely bead-associated growth recapitulates biofilm-like fungal organisation in human airways. In addition, approaches such as single-cell transcriptomics will be valuable for resolving cell-type–specific adaptations over time.

In summary, we present a reproducible and biologically relevant murine model of chronic pulmonary *Aspergillus* colonisation that bridges a critical gap between acute infection models and human airway disease. By capturing stable fungal persistence, restrained inflammation, epithelial remodelling, Th17-dominated adaptive immunity, and disease-specific pathology in CFTR-deficient hosts, this system provides a powerful framework for dissecting mechanisms of fungal tolerance, immune adaptation, and chronic lung remodelling. We anticipate that this model will facilitate future studies aimed at identifying therapeutic strategies that mitigate the pathological consequences of persistent *Aspergillus* colonisation without disrupting host–fungal equilibrium.

## Methods

### Mice

Wild-type C57BL/6J, CFTR-/+ and CFTR-/-mice were maintained under specific pathogen-free (SPF) conditions at University of Exeter Biological Services Unit. Animals were kept in individually ventilated cages with food and water ad libitum, on a 12-hour light/dark cycle (20–24 °C, 50–60 % humidity). Mice used in experiments were 6–8-week-old females randomly assigned to groups. All procedures were conducted under UK Home Office licence and approved by the University of Exeter Animal Welfare and Ethical Review Body (project license number: PP9965358).

### *Aspergillus fumigatus* airway colonisation

Conidia were cultured on minimal media agar plates at 37 °C for 5 days, harvested, washed twice in PBS containing 0.05% Tween-20 (Sigma), filtered through a 40 µm cell strainer, counted, and adjusted to 1 × 10^9^ conidia/mL. For bead preparation, molten 10% YPD–3.5% agar (5 mL, maintained at 62 °C) was mixed with an equal volume of concentrated conidia suspension or PBS containing 0.05% Tween-80 (Sigma) to generate empty controls. This mixture was added to 45 mL of prewarmed mineral oil (62 °C) and stirred vigorously at room temperature for 6 minutes, followed by cooling on crushed ice for 10 minutes to allow bead formation. Beads were collected by centrifugation (9,000 × g, 20 minutes, 4 °C), washed twice in PBS containing 0.05% Tween-80, and size-selected using sequential filtration through 100 µm and 70 µm strainers. The conidial load within beads was determined by homogenisation, plating on YPD agar supplemented with gentamicin (100 µg/mL) and vancomycin (10 µg/mL), and incubation at 37 °C for 48 hours. Following isoflurane anaesthesia, mice were intratracheally challenged with 50 µL of bead suspension containing 5 × 10^4^ CFU of *Aspergillus fumigatus* (Af293), or an equivalent volume of empty beads. At the indicated time points, lungs were harvested, homogenised in PBS, serially diluted, and plated on YPD agar for CFU enumeration after 48 hours at 37 °C. Clinical status and body weight were monitored daily.

### Assessment of *Aspergillus fumigatus* growth kinetics

Free or agar bead–embedded *A. fumigatus* conidia were cultured in MEM-α medium at 37 °C and 5% CO_2_. Fungal growth and morphogenesis were assessed by time-lapse Zeiss AxioObserver microscope at defined time points. Germination rates and hyphal length were quantified by tracking individual conidia using ZEISS ZEN 3.8 software.

### Epithelial cell activation

Human immortalised bronchial epithelial cells (HBEC) and human immortalised CF bronchial epithelial cells (CFBE41o^−^) were stimulated with sterile empty agar beads or left untreated as controls. After 24 hours, culture supernatants were collected and cytokine levels (IL-8 and IL-1β) were measured by ELISA following manufacturing instructions (R&D Systems).

### Tissue digestion and single-cell suspension

Lungs were dissociated using the lung dissociation kit following manufacturer’s instructions (Miltenyi Biotec). Spleen red blood cells were lysed with 1x RBC lysis buffer (BD PharmLyse, BD Biosciences). Bronchoalveolar lavage fluid (BALF) was collected following terminal anaesthesia. The trachea was surgically exposed and cannulated, and the lungs were lavaged by gently instilling and retrieving 1 mL of sterile PBS (in aliquots) through the tracheal cannula. Recovered fluid was pooled and kept on ice for downstream analysis. Cells were pelleted by centrifugation, and supernatants were stored at –80 °C for cytokine analysis. All suspensions were washed (400 *×g*, 5 minutes, 4 °C) and resuspended in complete RPMI (10% FCS, 1% penicillin/streptomycin, 50µM 2-mercaptoethanol) for tissue culture, or FACS buffer (PBS with 10% FCS, 2mM EDTA) for flow cytometry analysis.

### Flow cytometry

Single-cell suspensions were prepared in FACS buffer and stained with fixable viability dye (eFluor 780; eBioscience), washed, then stained with fluorochrome-conjugated monoclonal antibodies for same-day acquisition or fixed with 2 % PFA (Sigma). Conjugated monoclonal antibodies included: MHC-II-BUV496 (2G9, BD Biosciences), CD45-BUV496/PerCP-Cy5.5 (30-F11, BD Biosciences/BioLegend), CD4-BUV563/BV510 (RM4-5, RM4-4; BD Biosciences/BioLegend), CD3-AF594/eFluor 450 (500A2, BioLegend/eBioscience), B220-eFluor 450/AF700 (RA3-6B2, eBioscience/BioLegend), CD49b-eFluor 450 (DX5, eBioscience), CD11b-BUV395 (M1/70, BD Biosciences), Ly6G-eFluor450/SB550 (1A8, eBioscience/BioLegend), Ly6C-BV570 (HK1.4, BioLegend), F4/80-FITC/BV421 (BM8, BioLegend), CD11c-BV711 (HL3, BD Biosciences), NKp46-SB645 (29A1.4, eBioscience), NK1.1-BV605/AF647 (PK136, HP-3G10; BD Biosciences/BioLegend), Siglec-F-SB436 (1RNM44N, eBioscience), CX3CR1-BV785/PE (SA011F11, BioLegend), CD8-AF532 (RPA-T8, Thermo Fisher Scientific), CD64-AF647 (X54-5/7.1, BD Biosciences) and CD26-BUV737 (H194-112, BD Biosciences). For intracellular staining, IFN-γ-FITC (XMG1.2, BD Biosciences), IL-4-BV605 (11B11, BD Biosciences), IL-10-PE (JES5-16E3, BD Biosciences) and IL-17A-AF700 (TC11-18H10, BD Biosciences) were used. Intracellular staining employed the Transcription Factor Staining Buffer Set as per manufacturers instructions (eBioscience). Data were acquired on a Cytek Aurora and analysed with FlowJo v10 (BD Biosciences).

### Histology

Lungs were fixed in 10% neutral-buffered formalin, paraffin-embedded, sectioned at 4 µm, and stained with Periodic acid–Schiff (PAS) or Sirius Red staining. Tissue processing was performed by Microtechnical Services Ltd. (Exeter, UK). Images were acquired using an Olympus APX100 slide scanner, and quantification was carried out using QuPath software.

### Luminex multiplex cytokine analysis

Serum cytokines (CXCL1, MIP-1α, MIP-1β, TNF, MCP-1, RANTES, eotaxin, IL-12p40, IL-12p70, IL-6, IL-10, G-CSF, IFN-γ, IL-2, IL-4, IL-5, IL-17A) were measured using Bio-Plex Pro Mouse Cytokine kits (Bio-Rad) as per manufacturer’s instructions. Plates were read on a Luminex MAGPIX instrument and analysed with Bio-Plex Manager software.

### *In vitro* T cell restimulation

Splenocytes (2 × 10^6^ per well) were cultured in 96-well plates. For polyclonal stimulation, cells were treated with PMA (Sigma, 50 ng/ml) and ionomycin (500 ng/ml; Sigma) for 3 hours, followed by brefeldin A (10 µg/ml, BioLegend) for 3 hours. Cells were analysed by intracellular cytokine staining and flow cytometry.

### Statistical analysis

Statistical analyses were performed using GraphPad Prism 10. Fungal burden data were evaluated with Mann–Whitney U tests. For single comparisons involving one variable Student’s t-tests were used, while One-way ANOVA was applied for analyses involving multiple groups with Tukey’s multiple comparison test. Two-way ANOVA was employed when assessing the effect of two independent variables. Data are presented as mean ± standard error of the mean. unless otherwise specified. A P-value < 0.05 was considered statistically significant.

## Acknowledgements

We would like to acknowledge Prof. Gordon Brown for his support. We also gratefully acknowledge the Exeter Centre for Cytomics for assistance with flow cytometry analysis, and the University of Exeter Biological Services Unit for support with the animal experiments. This work was supported by the MRC Centre for Medical Mycology (MR/N006364/2 and MR/V033417/1) and the NIHR Exeter Biomedical Research Centre (NIHR203320). The views expressed are those of the authors and not necessarily those of the NIHR or the Department of Health and Social Care.

